# Reversal of visual feedback modulates somatosensory plasticity

**DOI:** 10.1101/2020.09.29.319350

**Authors:** Elana R. Goldenkoff, Heather R. McGregor, Joshua Mergos, Puyan Gholizadeh, John Bridenstine, Matt J.N. Brown, Michael Vesia

## Abstract

Reversed visual feedback during unimanual training increases transfer of skills to the opposite untrained hand and modulates plasticity in motor areas of the brain. However, it is unclear if unimanual training with reversed visual feedback also affects somatosensory areas. Here we manipulated visual input during unimanual training using left-right optical reversing spectacles and tested whether unimanual training with reversed vision modulates somatosensory cortical excitability to facilitate motor performance. Thirty participants practiced a unimanual ball-rotation task using the right hand with either left-right reversed vision (incongruent visual and somatosensory feedback) or direct vision (congruent feedback) of the moving hand. We estimated cortical excitability in primary somatosensory cortex (S1) before and after unimanual training by measuring somatosensory evoked potentials (SEPs). This was done by electrically stimulating the median nerve in the wrist while participants rested, and recording potentials over both hemispheres using electroencephalography. Performance of the ball-rotation task improved for both the right (trained) and left (untrained) hand after training across both direct and reversed vision conditions. Participants with direct vision of the right hand during training showed SEPs amplitudes increased bilaterally. In contrast, participants in the reversed visual condition showed attenuated SEPs following training. The results suggest that cortical suppression of S1 activity supports skilled motor performance after unimanual training with reversed vision, presumably by sensory gating of afferent signals from the movement. This finding provides insight into the mechanisms by which visual input interacts with the sensorimotor system and induces neuroplastic changes in S1 to support skilled motor performance.

## Introduction

Neural plasticity is the ability of the brain to adapt in response to changes in afferent input or efferent demand that often occurs with development, learning, injury and rehabilitation (Pascual-Leone et al., 2005; Fregni and Pascual-Leone, 2006; Dimyan and Cohen, 2011). This flexibility enables humans to readily modify behavior when learning new motor responses (Krakauer and Mazzoni, 2011). For instance, when visual and proprioceptive input of the hand and motor output are dissociated when wearing optical reversing prisms, individuals skillfully adapt to the altered visual feedback and are quickly able to perform accurate goal-directed movements despite the visuo-proprioceptive conflict (Stratton, 1897; Kohler, 1963; Sugita, 1996; Sekiyama et al., 2000; Marotta et al., 2005). In fact, there is mounting evidence indicating changes in the plasticity of both motor and sensory cortical areas during sensorimotor adaptation and motor learning (*for review, see* (Ostry and Gribble, 2016). Recent work has shown that changes in somatosensory cortical excitability occur during the early stages of motor skill learning in somatosensory cortical responses (Ohashi et al., 2019). Learning-induced enhancements in somatosensory evoked potentials (SEPs) in primary somatosensory cortex (S1) have been found following force-field adaptation, motor skill acquisition, and observational motor learning (Nasir et al., 2013; Andrew et al., 2015a; McGregor et al., 2016; O’Brien et al., 2020). It is known that parietal-frontal brain regions use sensory information to plan, execute, and adjust bodily actions (Gallivan and Culham, 2015). Given existing connections between primary motor and somatosensory cortices to premotor cortex, supplemental motor area, prefrontal cortex and parietal cortex (Graziano and Gross, 1998; Kakei et al., 2003; Dum and Strick, 2005; Fattori et al., 2015), this ability to adapt the bodily movement during the motor training task is likely associated with a distributed network of parietal-frontal brain areas involved in sensorimotor motor control (Andersen and Cui, 2009; Davare et al., 2011; Vesia and Crawford, 2012; Turella and Lingnau, 2014).

Unimanual training with reversed visual feedback has also been shown to improve motor performance in the opposite untrained hand – a process called intermanual transfer (Ruddy and Carson, 2013; Deconinck et al., 2014). Interestingly, this intermanual transfer of sensory and motor information has been linked to changes in neural plasticity in the motor, visual, and somatosensory systems (Hamzei et al., 2012; Lappchen et al., 2012; Nojima et al., 2012; Saleh et al., 2014; Ossmy and Mukamel, 2016; Saleh et al., 2017). It has been shown that unimanual training with reversed visual feedback enhances this intermanual transfer of motor skills and increases M1 excitability of the opposite untrained hand (Garry et al., 2005; Nojima et al., 2012; Yarossi et al., 2017; Jegatheeswaran et al., 2018). Given the interactions between motor commands and somatic perception in sensorimotor cortex during voluntary limb movement (Nelson, 1996; Umeda et al., 2019), it is conceivable that visual input could affect both motor and somatosensory excitability during motor learning (McGregor et al., 2016). Likewise, the effect of vision on tactile perception and changes to somatosensory cortical processing (Kennett et al., 2001; Ro et al., 2004; Haggard et al., 2007; Cardini et al., 2011) could modulate functional plasticity in sensorimotor regions during training.

Indeed, there is converging evidence indicating that the brain reduces proprioceptive inflow when performing tasks using reversed visual feedback in order to improve motor performance. For example, an individual with peripheral deafferentation performed significantly better on a mirror drawing task compared to healthy controls. The deafferented patient’s superior motor performance suggests that reducing proprioceptive inflow from the hand/limb can improve adaptation to visual perturbations in tasks involving visuo-proprioceptive conflict (Lajoie et al., 1992). Similarly, experiments involving healthy participants have shown that dampening somatosensory cortical excitability using repetitive transcranial magnetic stimulation prior to mirrored drawing results in improved performance outcomes (Balslev et al., 2004). Furthermore, visuomotor adaptation has been shown to suppress somatosensory input at the cortical level when visual and proprioceptive signals are discordant to support skilled movement (Bernier et al., 2009). Despite considerable evidence for a relationship between somatosensory gating (i.e., reduction in somatosensory inflow) and skilled motor performance during visuomotor adaptation, it remains unclear how the brain’s sensory gating during skilled motor control under reversed visual feedback might impact visuomotor adaptation and intermanual transfer.

The capacity of unimanual training and visual feedback reversal to induce motor plasticity and performance improvements in the untrained hand presents a promising avenue for interventions for individuals with unilateral motor impairments (e.g., mirror therapy) (Altschuler et al., 1999; Farthing et al., 2007; Ramachandran and Altschuler, 2009; Deconinck et al., 2014). However, our understanding of the mechanisms that lead to such sensorimotor brain and behavioral performance changes is limited. Here we tested the possibility that reversed visual feedback during unimanual training alters somatosensory plasticity and improves skilled motor control.

We tested the hypothesis that undergoing unimanual motor learning using reversed vision results in sensory gating to support skilled motor performance in both hands. In the current study, participants were trained to perform a ball-rotation task with their right hand. Participants underwent unimanual training using either direct vision (i.e., congruent visual and somatosensory inputs) or left-right (L-R) reversed vision of their right hand (i.e., visuo-proprioceptive conflict). Unimanual training with L-R reversed visual feedback gave the appearance that the training was performed using the left hand. We acquired SEPs to estimate S1 excitability changes associated with motor learning (Passmore et al., 2014; Macerollo et al., 2018). Before and after training, we used median nerve stimulation and electroencephalography (EEG) to elicit and record SEPs bilaterally. Similar to previous work (McGregor et al., 2016), we used the amplitude of the N20-P25 component of the SEP as an index of somatosensory excitability. After training with the right hand, participants performed the ball-rotation task using the untrained left-hand as an assessment of intermanual transfer. We predicted that unimanual training with the right hand would improve performance of the ball-rotation task in both the trained right-hand and the untrained left-hand, irrespective of visual condition. We predicted that the direct vision group would show increases in offline SEP amplitudes after training. In contrast, the L-R reversed vision group would exhibit sensory gating while training with conflicting sensory inputs, and as a result would not show offline SEP amplitude increases after training.

## Experimental Procedures

### Participants

A total of 30 healthy adults participated in the study, 15 of whom were assigned to a L-R reversed vision group (8 females, mean age: 24.9 ± 6.4 years, age range:18-38) and 15 who were assigned to a control direct vision group (9 females, mean age: 22.9 ± 4.8, age range:19-34). Data from one additional participant was collected but discarded from analyses because of poor signal to noise ratio in the EEG recordings. All participants were right-handed (Oldfield, 1971), had normal or corrected-to-normal vision, had no reported history of neurological or musculoskeletal disorders, and were naive to the behavioral task. Participants provided written informed consent to experimental procedures which were approved by the University of Michigan Institutional Review Board (HUM00144236).

### Experimental Protocol

The experimental design is shown in **Figure 1**. Participants were instructed to perform a skilled unimanual ball-rotation task with the left or right hand using two wooden balls (diameter = 3 cm), similar to a task used elsewhere (Nojima et al., 2012; Hoff et al., 2015; Rein et al., 2015). To familiarize participants with the task, all participants were first shown the same video of a tutor performing the clockwise (CW) ball-rotation task with the right hand from a top-down, first-person perspective. Participants then practiced the unimanual ball-rotation task with the right hand by performing CW rotations at their preferred speed with direct vision before testing. As a baseline measure, participants performed the unimanual task with the left hand by rotating the balls in the counterclockwise (CCW) direction around each other as quickly and accurately as possible for 30 s while looking directly at the left hand. We then acquired three 3-min-long SEP recordings first from the left hemisphere (right median nerve stimulation) and then from the right hemisphere (left median nerve stimulation) while participants rested both hands with their eyes closed. Next, participants were pseudo randomly assigned to the direct vision group or L-R reversed vision group for the unimanual training phase.

**Figure 1.**
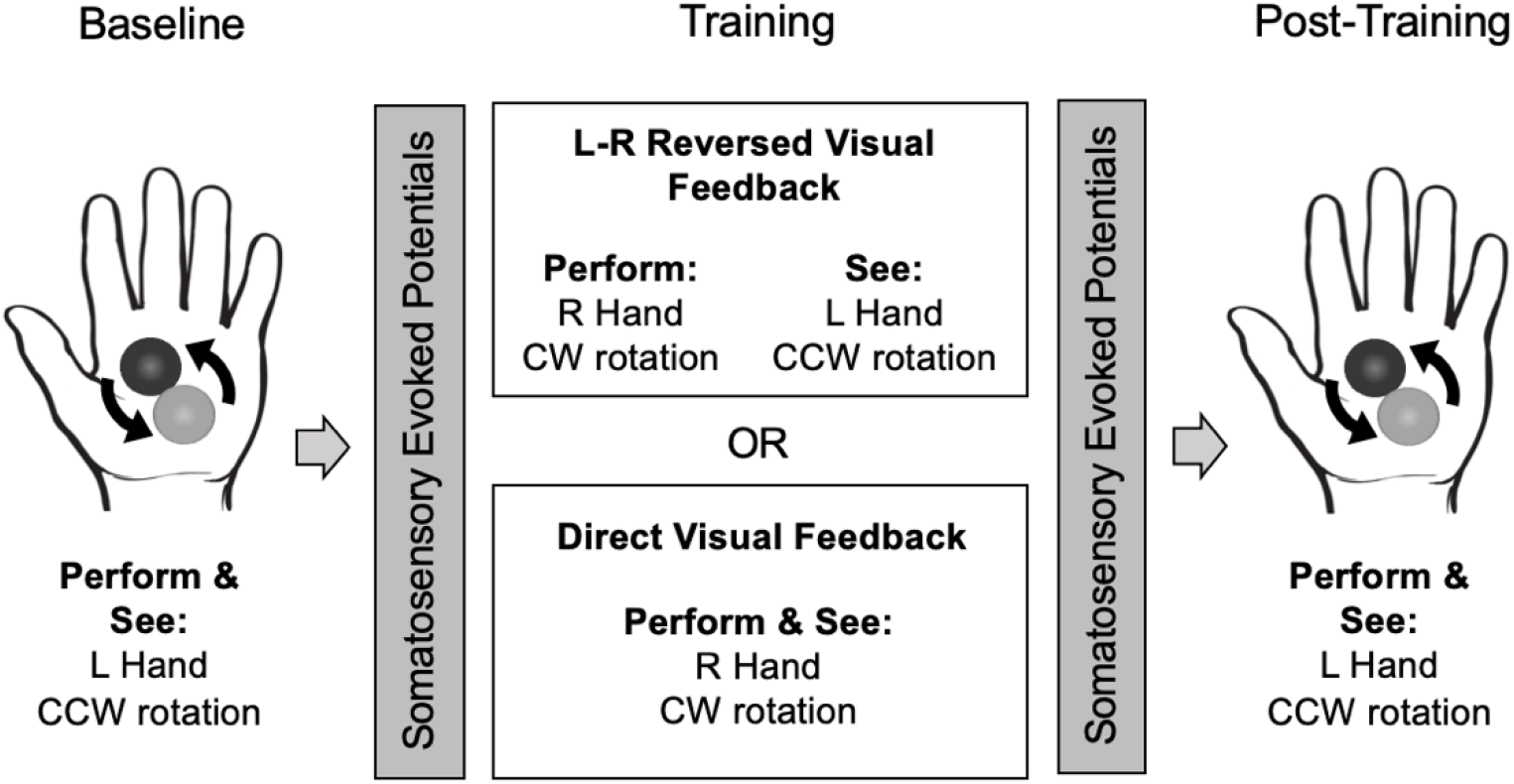
Experimental design. In the pre-training phase, all participants performed counterclockwise (CCW) ball-rotations with the left hand while looking directly at the left hand for 30 s. We then acquired somatosensory evoked potentials (SEPs) in primary somatosensory cortex (S1) from each hemisphere while the hands of the participant were relaxed. Next, participants were assigned to a training group which used either left-right reversed vision (incongruent visual and somatosensory feedback) or direct vision (congruent sensory feedback) of the right hand during training. During training, both groups performed ten 30-s blocks of clockwise (CW) ball-rotations with the right hand with 30-s rest periods between blocks. After training, SEPs were collected from each hemisphere to assess offline changes in SEP amplitudes in S1. To test for intermanual transfer, the number of CCW ball-rotations completed with the left hand in 30 s was assessed while participants directly viewed the moving left-hand.

During the unimanual training of the right hand, participants were instructed to perform the same ball-rotation task in a CW direction with the right hand. No nerve stimulation was applied during the motor task baseline, training, or post-training evaluation to avoid any confounds of increased cognitive effort and attention to the cutaneous sensation of nerve stimulation (Passmore et al., 2014). Participants performed the unimanual ball-rotation training with the right hand for a total of 10 blocks. Each block lasted 30 s, during which participants were instructed to perform CW rotations as quickly and accurately as possible. Participants were asked to fixate on the two wooden balls and pay attention to the moving hand while performing the motor task. Blocks were interleaved with 30 s periods of rest to avoid fatigue. The direct vision group performed the unimanual right-hand training while directly viewing the right (trained) hand, and thus received congruent visual and somatosensory feedback. The L-R reversed vision group viewed the right (trained) hand while wearing L-R optical reversing spectacles, giving the appearance that the left (inactive) hand was undergoing unimanual training in CCW direction. Consequently, participants in the L-R reversed vision group experienced incongruent visual and somatosensory feedback. The task presumably involves learning a visuomotor association, or mapping, between the visual input and appropriate reversed sensorimotor output (Sugita, 1996; Sekiyama et al., 2000; Krakauer and Mazzoni, 2011). Experimenters observed the participants performing the ball-rotation task and tallied the number of complete 360 degree rotations.

After the ten blocks of unimanual training with the right hand, we then acquired three 3-min-long SEP recordings first from the right hemisphere (left median nerve stimulation) and then from the left hemisphere (right median nerve stimulation) while participants rested both hands with their eyes closed. Finally, to assess intermanual transfer, participants were tested on the unimanual two-ball-rotation task in the CCW direction with the left hand for 30 seconds while looking directly at the left hand.

### Somatosensory evoked potential recordings

SEPs were acquired before and after right-hand unimanual training while participants closed their eyes and rested both hands. SEPs were elicited by bipolar stimulation electrodes placed along the left and right median nerves at the wrist with the cathode placed 2 cm proximal to the anode. Square wave pulses of 200 μs in duration were delivered at 2.79 Hz at the lowest intensity required to cause a muscle twitch in the thenar muscles (Cascade Elite, Cadwell, Kennewick, WA, USA). SEPs were recorded by Ag-AgCl scalp electrodes positioned at CP3 and CP4, overlaying the somatosensory cortical representation of the left and right hand, respectively. EEG signals were referenced to FPz according to the 10-20 system (Nuwer et al., 1994). A ground electrode was placed on the shoulder. We ensured that electrode impedances were below 5 kΩ. Data were sampled at 4 kHz and band-pass filtered online between 30 and 500 Hz. We acquired three 3-min-long SEP averages per hemisphere before and after righthand training. One SEP waveform was averaged from a total of 350 recordings that were time locked to the stimulus (0-50 ms) for further analyses. To examine S1 functional plasticity changes with skilled motor performance, we calculated the pre-training to post-training differences in the N20-P25 component of the SEP for each hemisphere. The N20-P25 complex is reproducible, generally unchanged by one’s cognitive state, and likely reflects the earliest cortical processing of afferent signals by S1 (Allison et al., 1991; Arthurs et al., 2004; Balzamo et al., 2004). Similar to previous work (McGregor et al., 2016), N20-P25 amplitudes were defined as the peak-to-peak measurement between the N20 and P25 components of each SEP, occurring approximately 18-25 ms after electrical stimulation of the median nerve at the wrist.

### Data analysis

We assessed changes in skilled motor performance during the right-hand unimanual training phase for each group. We first compared the number of complete CW ball-rotations performed by each group using the right hand during Block 1 versus Block 10 of unimanual training with a split-plot analysis of variance (ANOVA) using *Visual Condition* (2 levels: L-R reversed vision or direct vision) as a between-subject factor and *Block* (2 levels: Block 1 and Block 10) as a within-subject factor. To examine S1 functional plasticity changes with skilled motor performance, we calculated the pre-training to post-training differences in the N20-P25 component of the SEP in each hemisphere for each participant. A split-plot ANOVA was performed on the mean change in N20-P25 amplitude with *Visual Condition* (2 levels: L-R reversed vision or direct vision) as a between-subject factor and *Hemisphere* (2 levels: left or right) as a within-subject factor. We assessed intermanual transfer by comparing the number of CCW ball-rotations completed with the left hand during the pre-training versus post-training phase. We performed split-plot ANOVA on the number of CCW ball-rotations performed with the left hand using *Visual Condition* (2 levels: L-R reversed vision or direct vision) as a between-subject factor and *Time* (2 levels: pre-training or post-training) as a within-subject factor. We applied Greenhouse-Geiser correction when the assumption of sphericity was violated. We used a Bonferroni correction for *post hoc* comparisons. Partial η squared (η_p_^2^) values were computed as a measure of effect size, where appropriate. Cutoffs for effect sizes of ≥0.01, ≥0.06, and ≥0.14 are considered small, medium, and large, respectively (Cohen, 1992). Data in figures are presented as mean ± standard error. All statistical analyses were performed in IBM SPSS Statistics Version 26.0 (IBM Corp., Armonk, NY, USA). The threshold for significance was set at *p* ≤ 0.05 for all statistical tests.

## Results

### Motor skill performance of the right (trained) hand during training

Participants underwent unimanual training for ten 30 s blocks with the right hand. To assess the extent of skilled motor performance during right-hand unimanual training, we examined number of ball-rotations over the course of trial blocks for the direct vision (black, **Figure 2A**) and L-R reversed vision (grey, **Figure 2B**) conditions. A split-plot ANOVA conducted on the mean number of ball-rotations performed with the right-hand revealed a main effect of *Block* (F(1,28)= 41.3, *p*<0.001, η_p_^2^= 0.596), but no significant effect of *Visual Condition* (F(1,28)= 2.02, *p* =0.17, η_p_^2^=0.07), nor an effect of their interaction (F(1,28)= 1.97, *p* =0.17, η_p_^2^= 0.07). Post hoc comparisons with Bonferroni correction indicated that there was no significant difference between Prism and Direct conditions at training outset (Block 1 for direct vision versus Block 1 for L-R reversed vision, t(56) = 1.79, corrected *p*=0.16, *uncorrected p*=0.079). However, we found a performance improvement from Block 1 to Block 10, as evidenced by the greater number of ball-rotations completed with the right hand at the end of training phase, in both the direct visual condition (t(28)=3.56, *p*=0.0026) and the L-R reversed visual condition (t(28)=5.54, *p*<0.001). Therefore, the results suggest that participants in both groups demonstrated a similar level of improvement in motor skill performance with the right hand at the end of the training phase.

**Figure 2.**
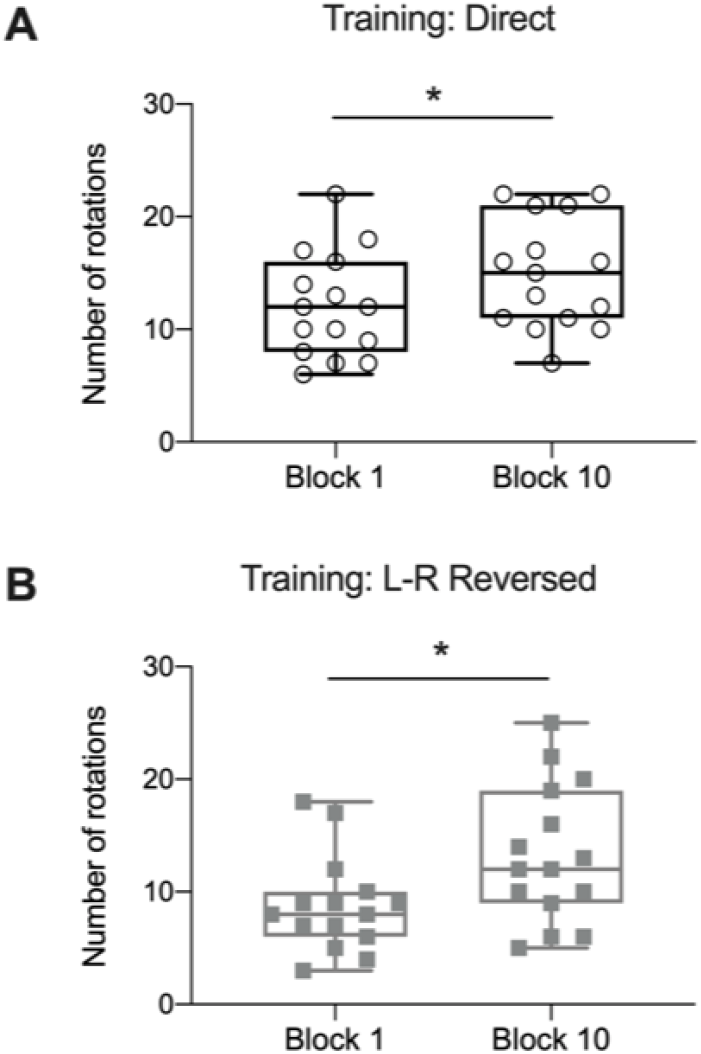
Performance of the right (trained) hand during training. Number of ball-rotations performed with the right hand for the first (Block 1) and last block (Block 10) of unimanual training in participants who either viewed the moving right-hand with direct vision (**A**, black, individual data points represented by open circles) or left-right (L-R) reversed vision (**B**, grey, individual data points represented by filled squares). Box represents median, 25th and 75th percentile and whiskers represent 5th and 95th percentile. *p≤0.05.

### Changes in somatosensory evoked potentials (SEPs) after training

We estimated offline S1 cortical excitability by comparing the change in the amplitude of the N20-P25 component of the SEP after participants performed a bout of right-hand unimanual training under direct vision or L-R reversed vision. A split-plot ANOVA conducted on the change in the N20-P25 component of the SEP revealed a main effect of *Visual Condition* (F(1,56)= 7.54, *p*= 0.008, η_p_^2^= 0.12). However, there was no significant effect of *Hemisphere* (F(1,56) = 0.37, *p* = 0.55, η_p_^2^= 0.01), nor their interaction (F(1,56) = 0.01, *p* = 0.92, η_p_^2^ > 0.01). Therefore, the direct vision group (black, open circles, **Figure 3**) showed larger increases in the mean changes in N20-P25 component amplitude across the hemispheres compared to the L-R reversed vision group (grey, filled squares, **Figure 3**). These results indicate that the differential effect on SEPs after unimanual training depends on the visual feedback of the hand during the bout of right-hand unimanual training (direct vision versus L-R reversal vision).

**Figure 3.**
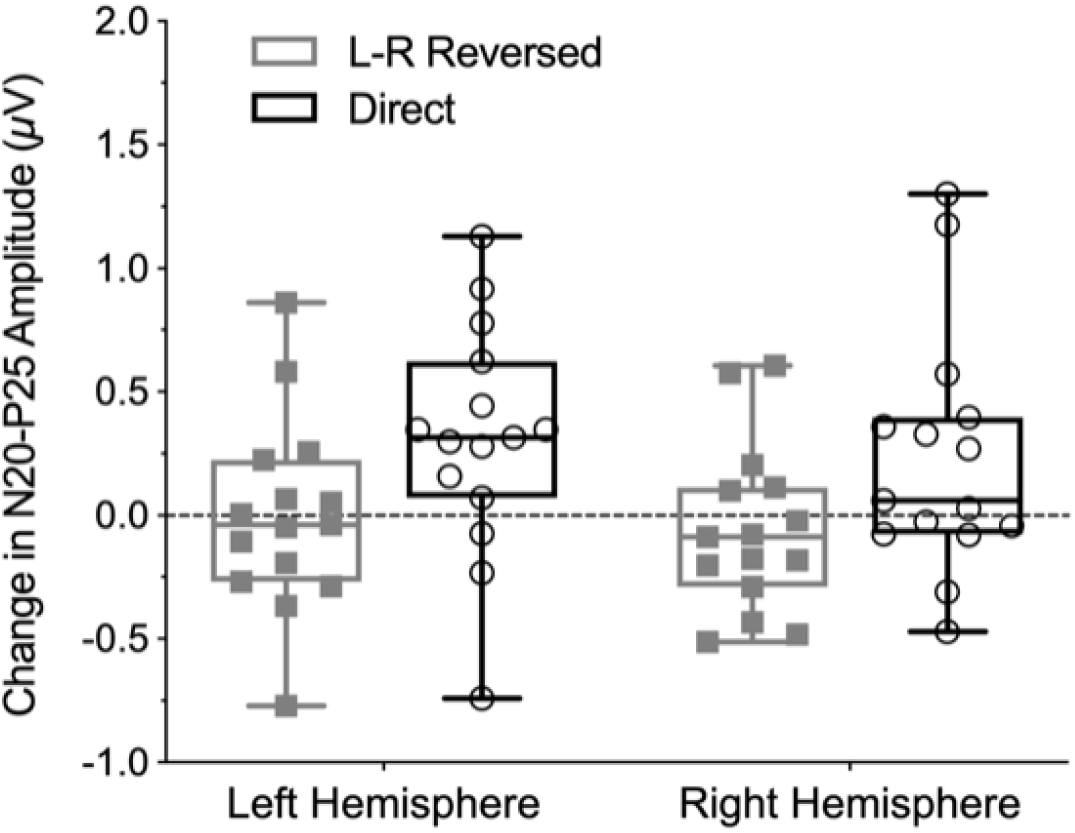
Mean change in somatosensory evoked potential (SEP) amplitudes after training. Mean change in N20-P25 component of the SEP from pre-to post-training for direct (black) or left-right (L-R) reversed vision (grey) group in each hemisphere. Change in the N20-P25 component amplitude for individual participants are represented by open circles and filled squares for direct and L-R vision respectively. Box represents median, 25th and 75th percentile and whiskers represent 5th and 95th percentile.

### Intermanual transfer to the left (untrained) hand after training

Before and after right-hand unimanual training, all participants performed CCW ball-rotations using the left hand under direct vision. We assessed intermanual transfer by examining changes in the number of ball-rotations performed with the left hand after undergoing unimanual training with the right hand under direct vision or L-R reversed vision. A split-plot ANOVA conducted on the mean number of ball-rotations with the untrained left-hand showed a main effect of *Time* (F(1,28) = 27.7, *p* < 0.001, η_p_^2^= 0.50), but no effect of *Visual Condition* (F(1,28) = 0.20, *p* = 0.66, η_p_^2^= 0.01), nor their interaction (F(1,56) = 3.08, *p* = 0.09, η_p_^2^= 0.10). *Post hoc* comparisons with Bonferroni correction found that both the direct vision group (t(28)=2.48, p=0.039, **Figure 4A**) and the L-R reversed visual group (t(28)=4.96, p<0.001, **Figure 4B**) completed a greater number of ball-rotations with the left hand after right-hand unimanual training compared to pre-training. The results suggest that unimanual training with the right hand enhanced motor-skill performance in the untrained left-hand, regardless of the visual feedback during the training phase.

**Figure 4.**
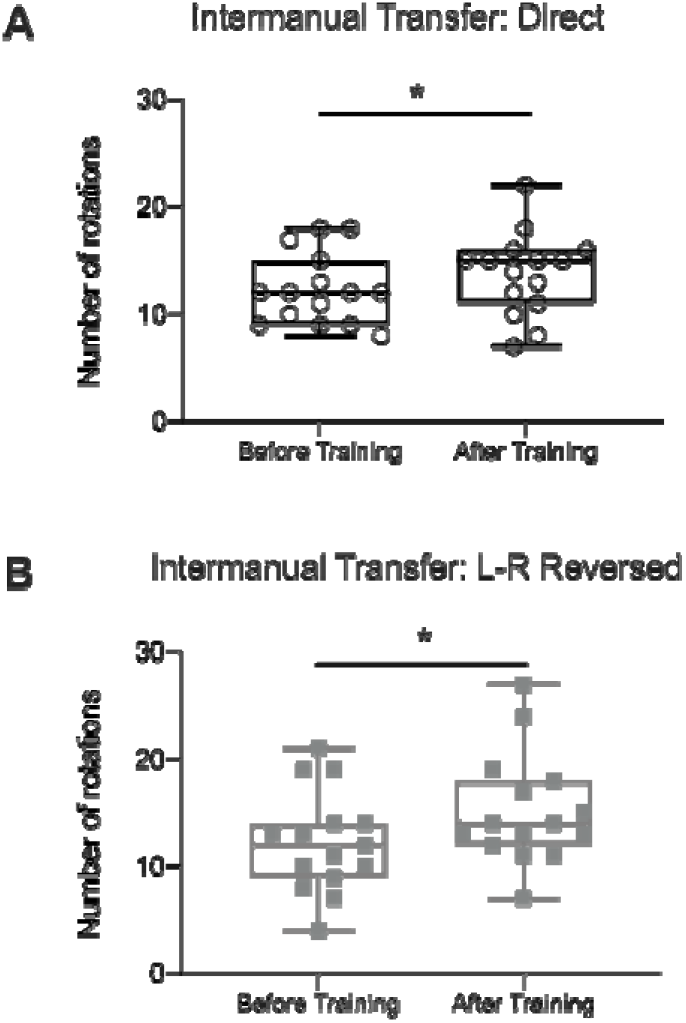
Intermanual transfer to the left (untrained) hand after training. **A**) Number of ball-rotations completed with the left hand in 30 s while participants directly viewed the left hand before and after right-hand unimanual training with either direct vision (**A**, black, individual data points represented by open circles) or left-right (L-R) reversed vision (**B**, grey, individual data points represented by filled squares). Box represents median, 25th and 75th percentile and whiskers represent 5th and 95th percentile. *p≤0.05.

## Discussion

The current study investigated the effects of reversed visual feedback during unimanual training on somatosensory excitability and skilled motor performance. SEPs measured before and after unimanual training showed that the direct vision group exhibited increases in offline SEP amplitudes in both the left and right hemispheres after training under congruent visual and proprioceptive inputs. This is consistent with the idea that plasticity in somatosensory areas coincides with learning-related changes to skilled motor performance (Vahdat et al., 2011; Nasir et al., 2013; Andrew et al., 2015a; 2015b; Sidarta et al., 2016; Ohashi et al., 2019) Critically, the observed bilateral increases in S1 excitability could be the result of interhemispheric interactions and transfer of sensory information between the left and right somatosensory cortices during the learning of the task; the resulting increases in SEPs recorded from S1 could be a product of Hebbian-like plasticity due to repeated bilateral sensorimotor activation during the unimanual movement. Thus, it is conceivable that the training-induced representation of the right hand could have modulated the functional hand representation in the opposite hemisphere. This idea is consistent with changes to the somatosensory system following conditioned SEP stimuli (Ragert et al., 2011), and bilateral S1 reorganization following amputation (Valyear et al., 2020) or temporary deafferentation with anesthesia (Werhahn et al., 2002). We further found that the reversed vision group, which performed the same task while using L-R optical reversing spectacles, did not show post-training changes in offline SEP amplitudes. These results indicate that functional plasticity in S1 after unimanual training was differentially modulated by visual feedback (direct versus L-R reversal). This suggests that reversal of visual feedback of the trained hand suppressed offline excitability increases in bilateral somatosensory cortices while still yielding improvements in skilled motor performance in both hands.

This finding aligns with previous evidence showing that cortical somatosensory activity is suppressed during experimental vision manipulations to reduce the perceptual conflict and facilitate the performance of motor skills when visual and proprioceptive signals are incongruent. Using EEG, Bernier and colleagues (2009) showed that early exposure to tracing with mirror-reversed vision attenuated the early arrival of somatosensory afferents to the contralateral S1 (N20-P27 complex), and covaried with motor performance, compared to tracing with direct vision (Bernier et al., 2009). It has been suggested that the gating of proprioceptive input during mismatched visual feedback may improve sensorimotor control to decrease intersensory noise in the motor system and minimize movement errors during adaptation. Interestingly, proprioceptively deafferented patients are less impaired in mirror drawing compared with healthy controls due to the lack of conflict between visual and proprioceptive signals (Lajoie et al., 1992), between a motor command and visual feedback of the resulting movement (Miall and Cole, 2007), or both. Similarly, reducing S1 cortical excitability with repetitive transcranial magnetic stimulation delivered to S1 improves motor performance on a mirror drawing task by presumably reducing the processing of proprioceptive input of the hand position (Balslev et al., 2004). We also found that using mirrored L-R reversed visual feedback during unimanual training suppresses post-training offline S1 excitability increases that occur following unimanual ball-rolling training with direct visual feedback.

Sensory reweighting may be a mechanism underlying the suppression of S1 excitability changes after training using L-R reversed vision. Through sensory weighting, the brain is thought to adjust its reliance on multisensory inputs such that reliable sensory inputs contribute more to performance while altered or unreliable inputs contribute less. The brain may downweight incoming somatosensory information in favor of visual input to optimize motor control (Touzalin-Chretien et al., 2010). When visual and proprioceptive feedback are incongruent during learning, it is possible that visual input of the hand was perceived as more reliable than the somatosensory inputs due to its higher spatial acuity (Sober and Sabes, 2005), resulting in decreased weighting on somatosensory inputs (Beauchamp et al., 2010). Evidence from behavioral studies indicates that vision can enhance tactile and proprioceptive perception (Kennett et al., 2001; Ro et al., 2004; Haggard et al., 2007), presumably by modulating activity of inhibitory intracortical circuits within the somatosensory cortex (Cardini et al., 2011). In this light, we recently found that the rubber hand illusion modulates the influences of somatosensory and parietal inputs to the motor cortex (Isayama et al., 2019). This is likely a consequence of the dominance of vision over proprioception which enhances a feeling of ownership of the seen body part (Botvinick and Cohen, 1998). Sensory reweighting is likely mediated by top-down gating of proprioceptive afferents from the periphery when adapting to novel visuo-motor mappings. Consistent with previous research, this work supports the idea that the brain relies on visual information over proprioception during visuo-proprioceptive incongruence to adjust to the novel visuo-motor mapping (Adamovich et al., 2009).

The effect of reversed vision is likely mediated by a large-scale reorganization in cortical areas within the sensorimotor network (Sugita, 1996; Sekiyama et al., 2000; Miyauchi et al., 2004; Deconinck et al., 2014). There is accumulating evidence from neuroimaging studies showing that mirrored visual feedback exerts a modulatory effect on the sensorimotor cortex through functionally interconnected parietal and frontal areas after training. For instance, unimanual mirror training induces neuroplastic changes such that functional connectivity increased between the premotor cortex and supplementary motor area corresponding to the untrained hand (Hamzei et al., 2012). Similarly, Tunik and colleagues have shown that mirrored visual feedback during unimanual training modulates activity within the sensorimotor representation of the contralateral, untrained hand (Tunik and Adamovich, 2009; Tunik et al., 2013; Saleh et al., 2014), and that this change is mediated by the contralateral parietal cortex (Saleh et al., 2017). A recent study also has provided evidence linking motor performance gains in the untrained hand with activity in the superior parietal lobule and the degree of coupling with motor and visual cortex with reversed visual feedback during unimanual training (Ossmy and Mukamel, 2016). These findings suggest that L-R reversed vision of the moving hand during unimanual training can influence plasticity throughout widespread areas in the sensorimotor system.

Given the substantial interconnections between visual, somatosensory, and motor cortical areas (Lewis and van Essen, 2000), afferent inputs about the hand position likely flow from primary and secondary somatosensory and visual areas to higher order cortical areas for processing. We surmise that the signals about the hand position converge in the parietal cortex where they are integrated for action-related processes (Andersen, 1997; Kalaska et al., 1997; Crawford et al., 2004; Culham and Valyear, 2006; Andersen and Cui, 2009; Vesia and Crawford, 2012; Fattori et al., 2015; Gallivan and Culham, 2015). We speculate that visual areas modulate cortical processing of somatosensory stimuli in S1 via recurrent projections from parietal areas to resolve the intersensory conflict about hand position during action control (Bolognini and Maravita, 2007; Limanowski and Blankenburg, 2016; 2017) before it is further processed in premotor areas (Graziano et al., 2000). The potent visual signal conveyed by the parietal cortex may be then sent to premotor and primary motor areas of the untrained hand directly or via S1 (Jones and Friedman, 1982). This signal, in turn, exerts a strong modulatory influence on motor excitability to support performance of motor skills (Garry et al., 2005; Fukumura et al., 2007; Funase et al., 2007; Merians et al., 2009; Carson and Ruddy, 2012; Hamzei et al., 2012; Lappchen et al., 2012; Nojima et al., 2012; Rein et al., 2015; Yarossi et al., 2017; Jegatheeswaran et al., 2018).

Given the compromised functional parieto-motor circuits following stroke and their association with the recovery of voluntary hand movements, a better understanding of the effects of unimanual training with L-R reversed vision on the motor system could play an important role in developing neuromodulatory therapies (Grefkes et al., 2010; Grefkes and Fink, 2014; Grefkes and Ward, 2014; Schulz et al., 2015; 2016). Although our findings may prove clinically useful, caution should be taken when drawing conclusions from the effect of a single bout of practice on functional plasticity in the somatosensory cortex. Subsequent studies also should include additional electrodes placed over other cortical areas as a control to determine the specificity of the effect. Another methodological limitation of our study is that we did not record changes in evoked potentials at later latencies (i.e., at 50 ms post-stimulus) that would have provided further knowledge of the mechanisms contributing to the change in S1 plasticity. As a result, we cannot rule out the possibility that attentional and cognitive modulations from parieto-frontal areas can modulate visuo-proprioceptive gain which in turn adaptively adjusts motor output. This adjustment may resolve intersensory conflict which has been shown to contribute to the suppression of S1 activity (Fink et al., 1999; Limanowski and Friston, 2019). We also did not manipulate somatosensory brain areas to establish if they play a causal role. Future work should investigate the cumulative effect of repeated unimanual training with reversed vision (i.e., days or weeks) to better understand the persistent changes in training-induced plasticity on motor function. Given our moderate sample size, a critical issue relates to whether there is a relationship between changes in SEPs and measures of behavioral improvement. Although one would expect this relationship to be present based on recent findings where participants with larger changes in N20-P25 magnitudes showed superior motor performance (Ohashi et al., 2019), it will be important for future studies to investigate this possibility with a larger sample size. Finally, we cannot speak to the duration of the somatosensory attenuation because we only acquired one measurement immediately after training rather than at multiple time points (e.g., 15 mins, 30 mins, 1 hr post-training).

In summary, our results support the idea that unimanual training with reversed vision suppresses learning-induced S1 activity changes while still facilitating improvement in skilled motor performance. This may occur via sensory gating of somatosensory or proprioceptive inputs from the movement. This knowledge and the ability to quantify changes to somatosensory plasticity that accompany unimanual training with reversed vision may provide a better understanding of the effects of rehabilitation in patients with unilateral orthopedic and neurological impairments (Altschuler et al., 1999; Farthing et al., 2007; Ramachandran and Altschuler, 2009; Deconinck et al., 2014; Zult et al., 2015).

## Acknowledgements

This research was supported by a pilot grant from the School of Kinesiology at the University of Michigan. HRM was supported by a NSERC postdoctoral fellowship.

